# Auditory responses evoked by optical stimulation on the optogenetic-infected cochlear neurons in the guinea pigs

**DOI:** 10.1101/2020.11.14.375519

**Authors:** Chen Liu, Shu Fang, Da-xiong Ding, Han-dai Qin, Shuo-long Yuan, Ping Lv, Yu Ning

## Abstract

Cochlear implants (CIs) are by far the optimal option to partially restore hearing for the patients of sensorineural hearing impairment (HI) by electrically stimulating spiral ganglion neurons (SGNs). However, wide current spread from each electrode constitute an interface which restricts precision and quality of the electrical CIs. Recently, optogenetic stimulation of the cochlea has been proved as a more optimized approach via adeno-associated virus (AAV) carrying the gene encoding the light-sensitive channelrhodopsin-2. Here, we focus on summarizing recent work on stable and accurate ChR2 expression and compare the electrophysiological recording of optogenetic and acoustic stimulation in adult guinea pigs. Light stimulation generated auditory responses that was similar to that of acoustic stimulation. Moreover, normal hearing adult guinea pigs responded with a rise in amplitudes with increasing light intensity. In conclusion, optogenetic cochlear stimulation achieved good spectral selectivity of artificial sound encoding in a new adult rodent model, suggesting that the capabilities of optogenetics might be applied to improve cochlear implants in the future.

## Introduction

Hearing impairment is widespread and found all over the word. An estimated 466 million people worldwide – 6.1% of the population – have disabling hearing loss, and this number is expected to rise^[1]^. Cochlear implants (CIs) have been the most effective restoration of hearing for the deaf to allow them to hear speech and sound^[2]^.Despite its success, shortcomings of CIs are also apparent; for example, there are up to about 20 frequency channels for the patients with cochlear implants compared to the normal physiological frequency channels^[3]^. Electric stimulation limits the spectral resolutions that ultimately result in limited perception of acoustic signals, such as speech, especially in noisy environments^[4]^.Optogenetics, combining genetic and optical techniques, can specifically target the confined spiral ganglion neurons (SGNs)with high precision^[5–7]^, reducing the limitation of channel cross-talk from wide spread of current around each electrode contact^[8]^. Genetic manipulation and expression of transgenes encoding alien proteins have been proven not to bring risks for adverse effects such as immune responses or proliferation, neuronal toxicity or loss^[9]^. Transgenic mice^[10]^ and rats^[11]^ or the ones that use trans-uterine virus injections into the otocyst of available embryos are proved to be available to activate of the auditory pathway^[12]^.However, there is a long way to achieve the real clinical translation. Here, the goal was to establish the adult animal models of optogenetics which express opsins in auditory neurons safely and effectively. AAVs are good candidates because they were successfully used to transduce murine SGNs^[13, 14]^. Therefore, Virus (AAV)–mediated gene transfer is a standard approach in optogenetics^[15]^. Although adult Mongolian gerbils have been established with AAV-mediated expression of CatCh in SGNs^[16]^, the success rate remains to enhan ce on their way to potential clinical translation further. Concerning about the postnatal approach for manipulating SGNs throughout, the whole cochlear must be raised biologically safe and take efficient protocols for genetic manipulation of SGNs. Meanwhile, identify the expression of ChR2(H134R) on the adult guinea pig at both molecular and cellular level. Using the electrophysiological way to analysis the relationship between the laser intensity and the potential of inner ear.

## Materials and Methods

### Animal preparations

Healthy adult guinea pigs (200–250 g) of either gender were used for the experiments. All of the animals were bought from Vital River Laboratory Animal Technology Co. Ltd. (Beijing, China). For each surgery, guinea pigs were anesthetized with an intraperitoneal injection of pentobarbital sodium 40mg/1kg diluted in 0.9% sterile NaCl solution before surgery. The depth of anesthesia was monitored by the hind limb withdrawal reflex and adjusted accordingly. During all experiments, animals were placed on a heating plate, and body temperature was maintained at 37 °C. All procedures in this study were approved by the Institutional Animal Care and Use Committee of the Chinese PLA General Hospital. Animals should be humanely euthanized at the end of the procedure. Suitable methods of euthanasia may adopt cervical dislocation.

### Vector preparations

Viral vectors used in this study was AAV-2/8.AAV2/8 carrying ChR2(H134R) linked to the reporter protein enhanced yellow fluorescent protein (eYFP) under control of the human synapsin promoter for injections into the cochlea of adult guinea pigs. In order to transduce in vivo, we adopted 1.10E+13 v.g./ml as a standard concentration of viral inoculum and injected about 5μl of AAV-ChR2(H134R) suspension into the cochlea.

### Injection of vector

Injections were performed by using micropipettes pulled from pipettes after heating. The micropipettes were connected to a microinjector under control of micro-injection pum. About 5 μl of virus suspension were injected directly into the right inner ear of adult guinea pigs through the round window membrane. Make an incision behind the right ear, dissect the muscles and connective tissue covering the bulla tympanica and perform an ostomy on the round window membrane. After injection, gelatin sponge was covered on the round window membrane. Muscles and connective tissue were repositioned and the skin was sutured. Animals were allowed to recover for at 4-6 weeks after surgery before continuing experiments.

### Surgical Procedure

1. Check the signs of the paw withdrawal reflex after 15min anesthesia induction, and proceed only if the reflex is absent.
2. Carefully shave the retroauricular area in preparation of the incision. Use a retroauricular approach to reach the bulla (air-containing middle ear cavity).
3. With a mucous membrane knife carefully dissect and remove part of the muscles covering the bulla.
4. Identify the area as a bony structure with rough granules on the bulla surface that indicates the position inserting the tympanum. Punching with the miro electric drill on the place above the round window. Then, the round window and promontorium of the cochlea is exposed.
5. Inserting the miniature catheter into the round window. Inject the virus into inner ear in 1min with micro-injection pum, then cover the round window membrane with gelatin sponge.
6. For a round window insertion of an optical fiber implant to scala tympani, carefully enlarge the incision above the round window with micro polishing drill at low speed.

### Optical Stimulation

1. For laser stimulation, use a 470 nm continuous wave laser. Couple the optical fiber to the 470nm blue laser source that can ideally offer fast control of power.
2. For intracochlear stimulation, insert the fiber through the round window into the scala tympani. Then use 5ms laser pulses of 3.70 mW (power of fiber output) to elicit oCAP. The large amplitude of oCAP enables the experimenter to observe oCAP evoked by optical stimuli on the oscilloscope for optimizing the position and orientation of the emitter (fiber). The read-out represents the the amplitude of oCAP.
3. Adjust the position of the fiber. When the waveform of oCAP was stable, fix the fiber. NOTE: It is normal to observe some liquid coming out from the cochlea or exfusing from the tissues, which affect the amplitude. It is necessary to clean it in time in order to avoid the bad influence, and it is advisable to dry up the region using fine cotton or sponge.

### Optical compound action potential (oCAP) recording

Guinea pigs were anesthetized by pentobarbital sodium (40mg/1kg, ip).Body temperature was maintained at 37–38°C by placing anesthetized guinea pigs on a heating plate. (surgery procedure). The silver electrode was placed on the round window membrane and the reference electrode was placed in the muscles around the ear. .,/n

Optical compound action potential (oCAP) was recorded in a double-wall sound isolated room by use of a Tucker-Davis ABR workstation (Tucker-Davis Tech. Alachua, FL, USA). Sample at a rate of 3.1times/sec and filter (300-3,000 Hz). Visualize the different potential on an oscilloscope ideally near the recording setup for optimizing the position of the optical stimulator. Use signal averaging (100 times) and store the data on a computer for offline analysis.

### Immunofluorescent staining and confocal microscopy

Drop 0.25% Triton X-100 solution for 30min at room temperature on the slices; rinse with PBS for 3 times, 5 minutes each time; Add the blocking solution diluted primary resistance (mouse resistance to NF 1:400 and rabbit resistance to M-cherry 1: :400), and then incubate at room temperature for 2h. Suck up the primary resistance and rinse it with PBS for 3 times, 5min each; Add the secondary resistance diluted with blocking solution (sheep anti-mouse Alexa-488 1:400, sheep anti-rabbit Alexa-568 1:400), and incubate for 1 hour at room temperature and avoid light, and rinse with PBS for 3 times, 5 minutes each time; Add DAPI sealant to the tissue. Leica SP8 laser confocal fluorescence microscope was used to observe the antibody expression of the specimens under 20X and 40X and the images were collected. Photoshop CS6 software was used for post-processing.

### Statistical analysis

Data were plotted by SigmaPlot and statistically analyzed by SPSS v25.0 (SPSS Inc. Chicago, IL). Data were expressed as mean ± s.e.m. The two-tailed Student’s t test or one-way ANOVA was used. The threshold for significance was α = 0.05. Bonferroni *post hoc* test was used in ANOVA.

## Results

### Optogenetic manipulation of the spiral ganglion in adult guinea pigs via round window injection

We used AAV2/8 carrying ChR2 (H134R) linked to the reporter protein-enhanced yellow fluorescent protein (eYFP) under control of the human synapsin promoter for miroinjections through the round window into the cochlea of adult guinea pigs (8 to 15 weeks of age). We adapted a retroauricular approach to open the bulla and approached the round window using a fine dental drill. Through the round window to inject 3 to 5 ul of AAV-ChR2 (H134R) suspension (Fig. 1, A to C). We visualized the expression of ChR2 (H134R) by using various verification routes in adult guinea pigs. ChR2 (H134R) showed an efficient and sufficient expression on the plasma membrane of SGNs.

**Fig. 1.**
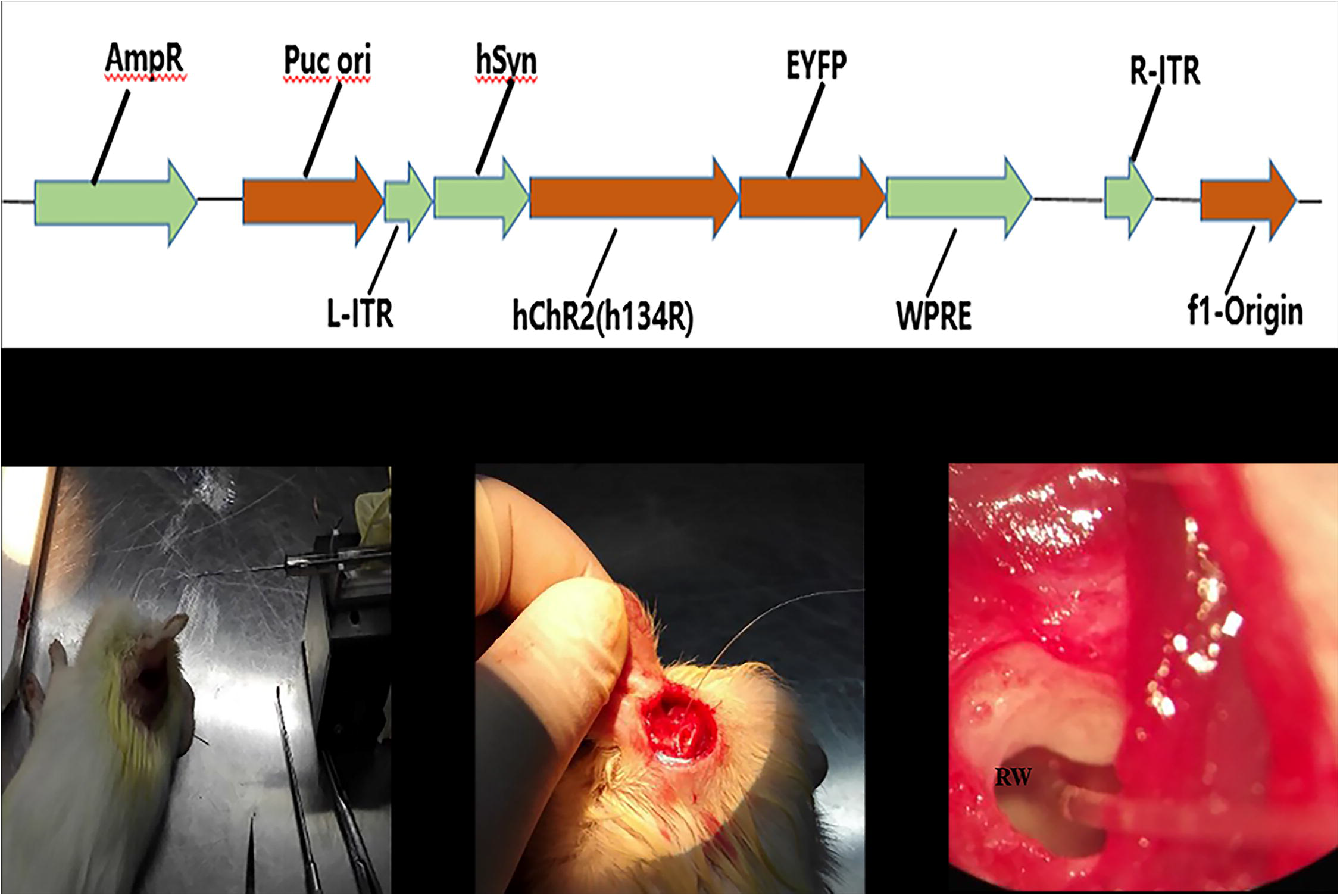
AAV2/8-ChR2(H134R)-mediated optogenetic manipulation of cochlear SGNs in adult guinea pig. (A) Scheme of the AAV2/8 construct used to transduce SGNs with ChR2(H134R). hSyn, human synapsin promoter. (B,C,D) Depiction of retroauricular incision and the picture of the retroauricular approach to the round window of the guinea pig (RW, round window). Images showing inserting the catheter to the round window, connect the catheter to the microsyringe and use a micro-injection pum to inject the virus into inner ear.

### Optogenetic Expression of ChR2 in the Spiral Ganglions

4 to 6 weeks later, after injection, the expression of ChR2 (H134R) - eYFP of SGNs were analyzed by confocal microscopy of immunolabeled cochlea cryosections and RT-PCR. Both of the two methods reliably and effectively proved the expression of ChR2 (H134R)-eYFP, which was limited to SGNs of the injected ear compared to the noninjected side. Immunohistochemistry of cochlear cryosections showed accurate labeling in SGNs (Fig. 2). By analyzing the colocalization of anti–NF200 and eYFP (with ChR2) with software of Imari, we further confirmed the expression of ChR2 is on the membrane of somata and neurites (Fig. 3). The yellow regions represent the overlap of NF200 and eYFP, which are confirmed the expression of ChR2 on the membrane of SGNs (Fig. 3).

**Fig. 2.**
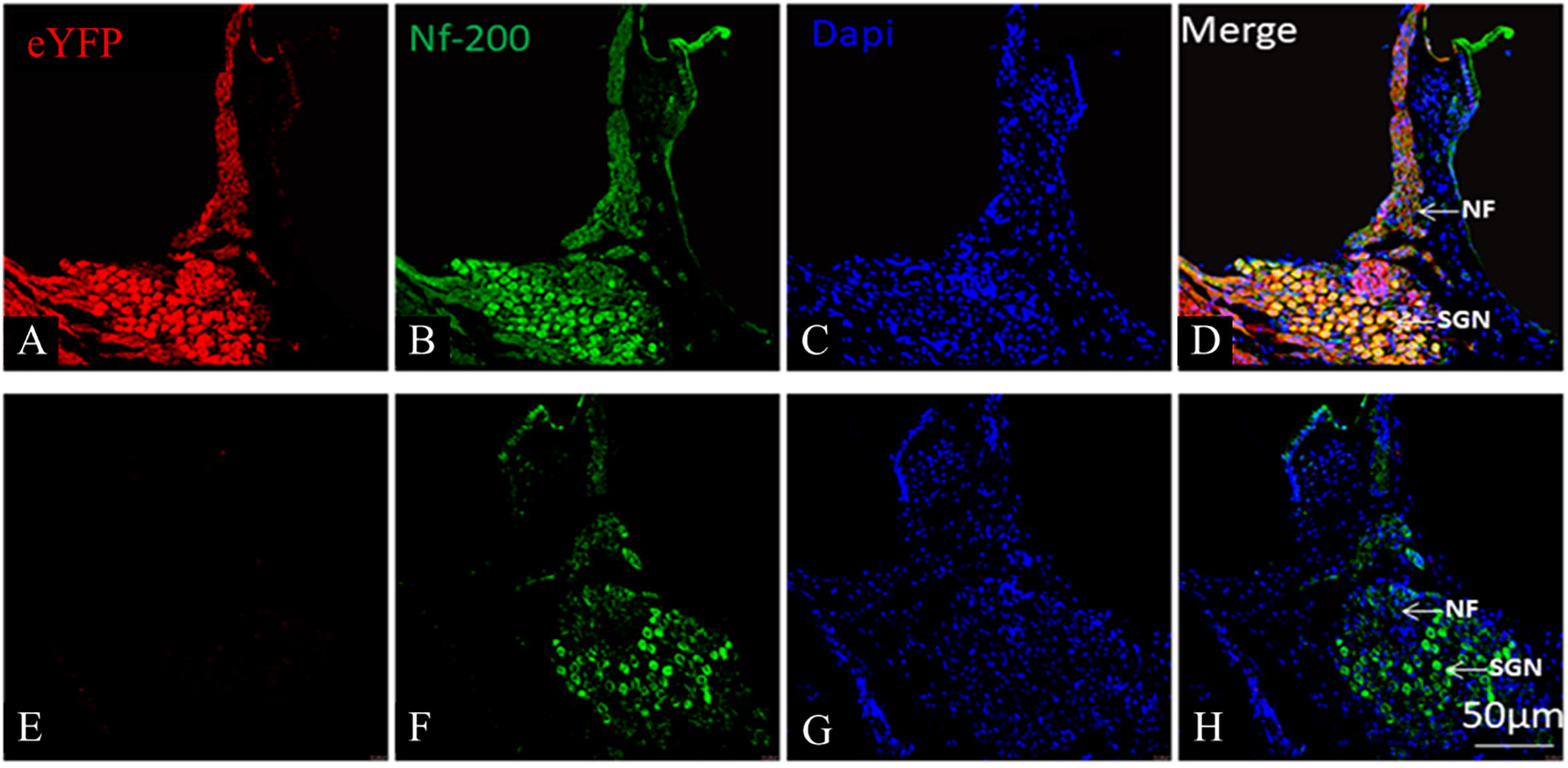
Confocal images of immunolabeled cochlear cryosections (representative section of a spiral ganglion) of an AAV-ChR2(H134R)–injected adult guinea pig. eYFP (green) marks transduced SGNs, DAPI (blue) marks cell nucleus and neurofilament-200 generically marks SGNs (red). Scale bars, 50μm. It should be noted that the apparent abundance of ChR2(H134R) demonstrates a clear membrane expression. Magnification of A~D showing ChR-2 expression in SGNS, E~H is control under the same immunostain.

**Fig. 3.**
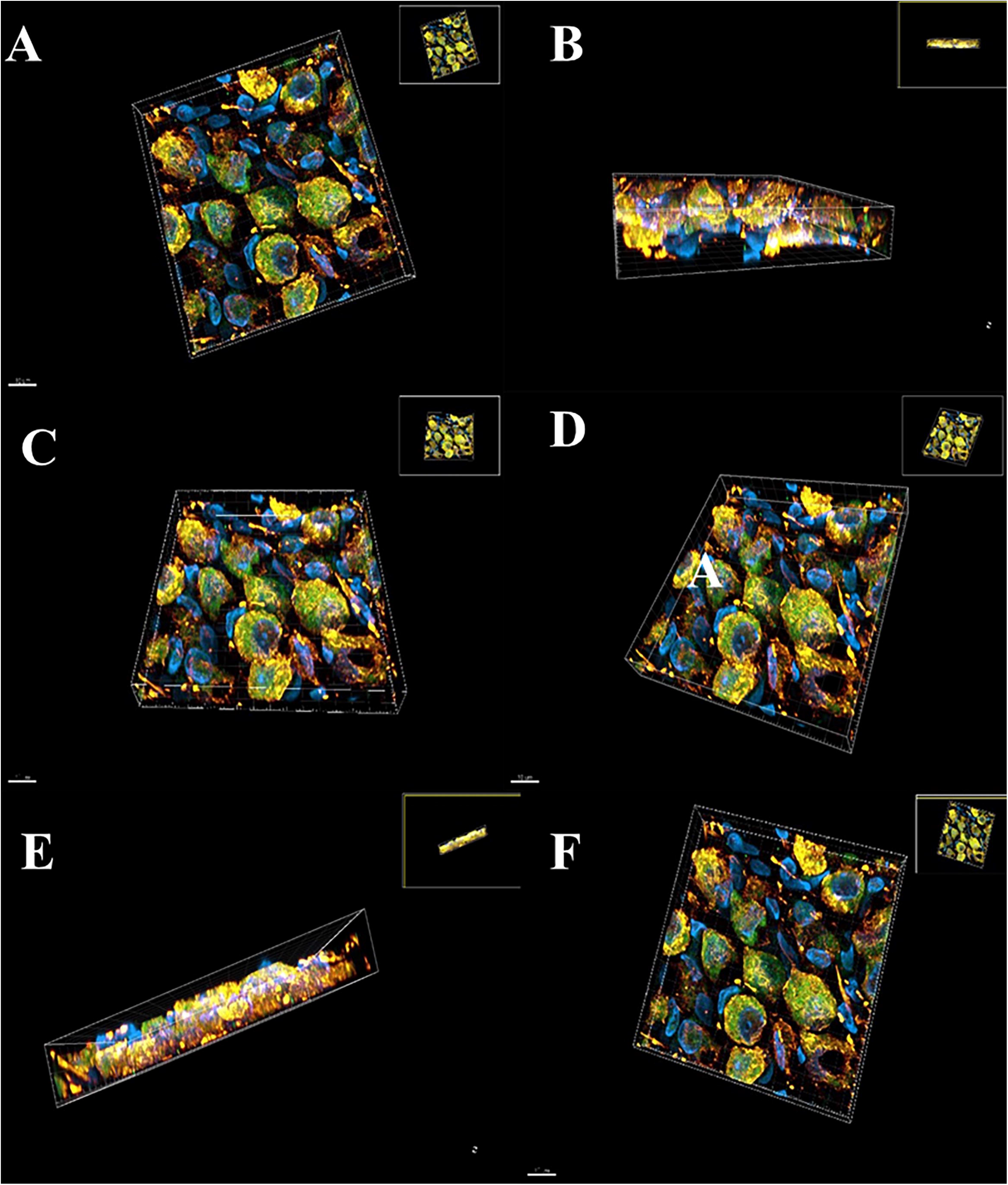
The colocalization of anti–NF200 (red) and eYFP with ChR2 (geen) from every angle of SGNS(A-F. The yellow represent the overlap of NF200 and eYFP, which are confirmed the expression of ChR2 on the membrane of SGNs

### The RNA for ChR2 (H134R)-eYFP is up-regulated by injection in SGNs

Before assessing functional expression of ChR2 from SGNs, we first observed a robust fluorescent signal of neuronal cell bodies and fibers in the SGNs (Figure 2, 3). To verify the expression of ChR2 on SGNs, we performed TaqMan RT-PCR for ChR2 and eYFP. Indeed, the neurons of auditory expressed ChR2 and eYFP. Primers for the genes that encode for ChR2 (H134R), eYFP, actin (Actb) were designed using Clone Manager software (Sci Ed Software). We performed quantitative analysis of mRNAs that encode for ChR2 (H134R), eYFP, actin (Actb) using qPCR on SGNs from adult guinea pig (Figure 4A). This analysis revealed that ChR2 (H134R) and eYFP mRNAs expressed in the SGNs of guinea pigs compared to the control. This analysis revealed that ChR2 (H134R) and eYFP mRNA is higher in injected guinea pig compared with non-injected ones (Figure 4B), whereas levels of actin (Actb) mRNA were not essentially different from the two groups. (Figure 4B).

**Fig. 4.**
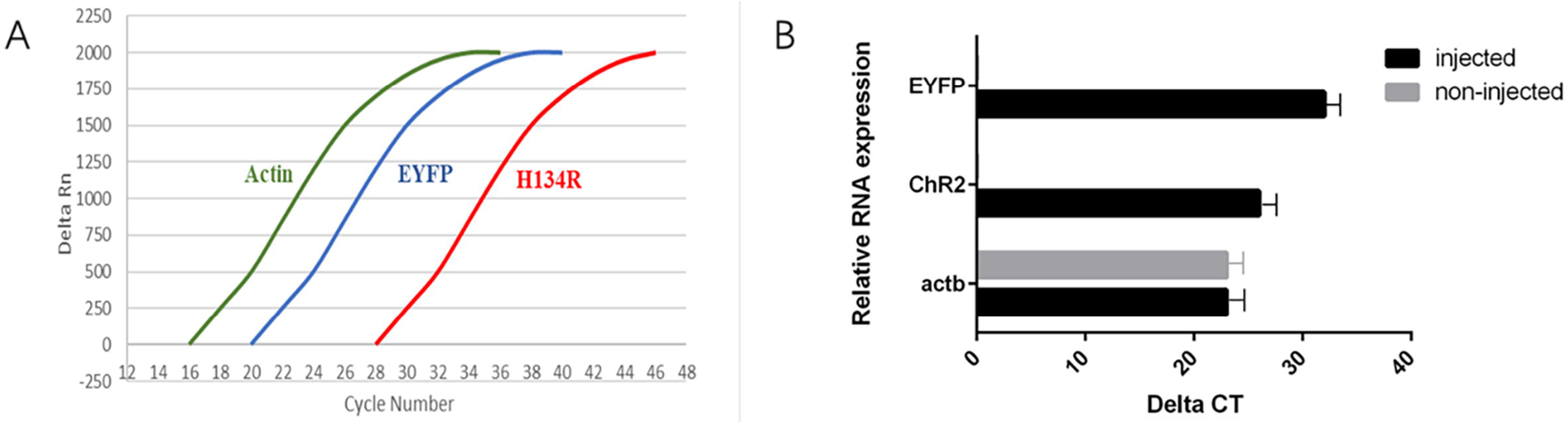
ChR2 (H134R),EYFP, actin (Actb) expression in SGNs. A, qPCR assay with amplification curves for ChR2 (H134R),EYFP, actin (Actb). Cycle number of injected guinea pigs was plotted against the normalized fluorescence intensity (Rn) to visualize the PCR amplification. Curves depict the expression of ChR2 (H134R) and EYFP in the SGNS of injected guinea pigs. B, Bar graph summarizing RNA expression of ChR2 (H134R),EYFP and actin (Actb) from SGNS of injected guinea comparing to the non-injuected ones. Animal numbers are indicated (on average). actin (Actb) was detected in both injected and non-injected guinea pigs,however,ChR2 (H134R) and EYFP RNA in the SGNs of the experimental group was detectable.

### Activation of the auditory pathway by optical stimulation

Because ChR2 expression was restricted to SGNs, the optogenetically evoked auditory activity can be safely attributed to SGNs. We respectively tested the activation of the auditory pathway by acoustic and optogenetic stimulation using round window recordings of neuronal population responses (Compound Action Potential, CAP : the sum of the action potentials produced by thousands of nerve fibers, acoustic CAP(aCAP): averaging 1,024 trials and oCAP: averaging 100 trials, respectively). Then, characterizing the laser-induced activation of the auditory pathway, we recorded oCAPs by using fiber-coupled laser stimulation with different blue light stimulation strategies. Due to the middle ear surgery and virus-injection, the acoustic auditory brainstem responses(ABR) or CAP were mildly compromised (Figure 5A), as illustrated by the reduced amplitudes of the suprathreshold aABR and an elevation of the auditory threshold and latent period compared to the normal saline control (11–12dB, n = 20, P <0.01 by a paired Student’s t test) (Figure 5B,C). oCAP differed from aCAP (evoked by 40 dB clicks) in waveform, number of waves, and amplitude (Figure 5E). Quantitative comparison of aCAP (stimulus rate:21times/sec; duration: 10 ms; intensity, 40dB dB SPL) from noninjected guinea pigs and oCAP data with the highest laser intensity (stimulus rate:3.1times/sec; duration:10ms; intensity, 5.80mW) showed similar amplitudes and nearly equivalent latency (oCAP evoked by 5.80mW blue light: amplitude 0.78μV, latent period 2.71ms; aCAP evoked by 40dB SPL click: amplitude 0.70μV, latent period 2.59ms). This may reflect a higher synchrony of more SGNs induced by optogenetic versus acoustic stimulation. We did detect oCAP response to the different stimulation intensity of the laser transcochlear irradiance in the adult guinea pig. Comparing the relationship between amplitudes and laser intensities, We conclude that the amplitudes increased with laser intensity rising (Figure 5F). Nevertheless, when laser was projected into their control cochlea, any action potential was evoked. There were slight differences in amplitude, waveform and latency among each experimental animals, but were reproducible within the same recording. Together, these results suggest that the ChR2 (H134R) optogenetic coding achieves a temporal fidelity similar to that of acoustic coding on the SGNs level.

**Fig. 5.**
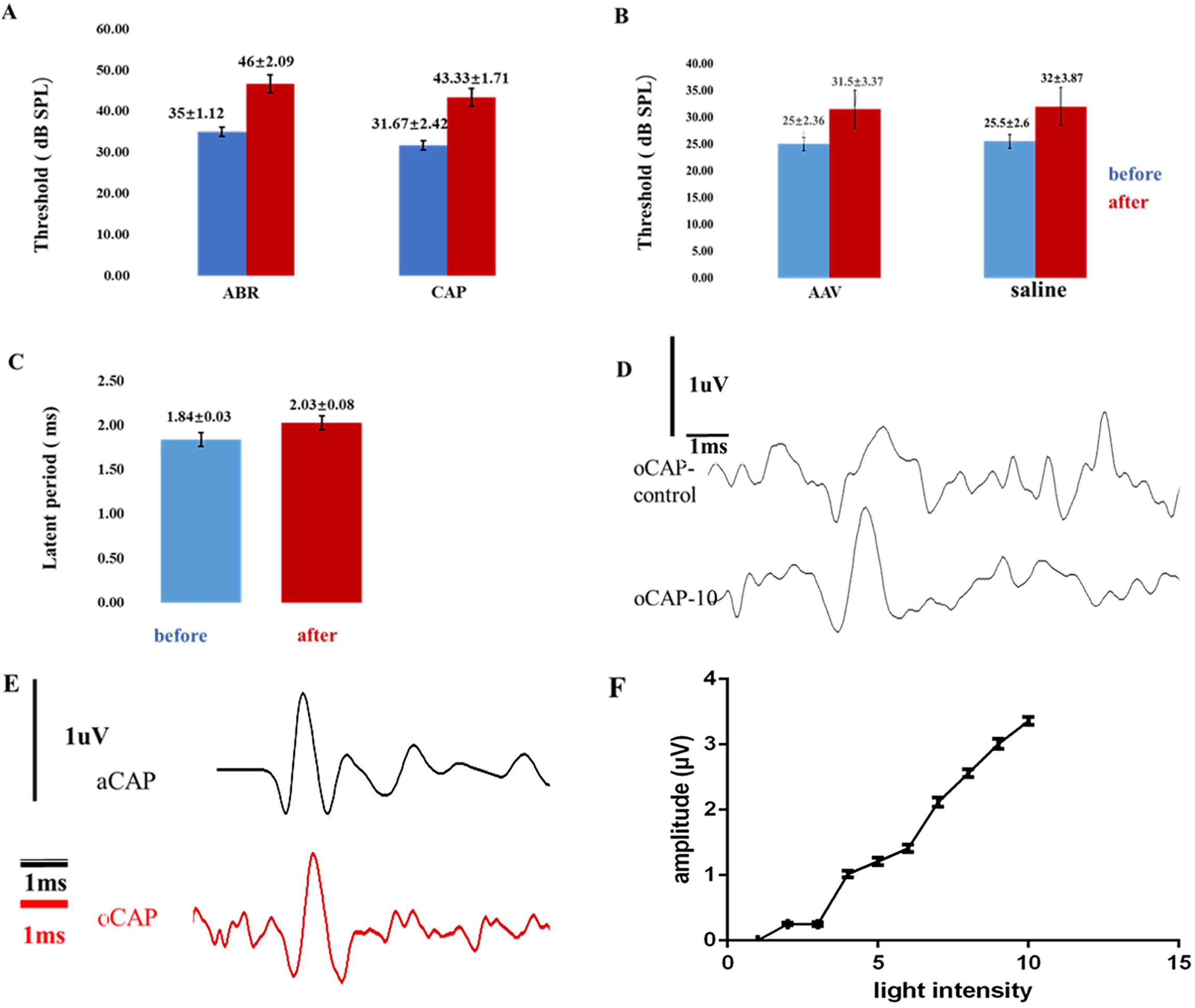
Optical activation of the auditory pathway in the injected guinea pig.A, Average aABRs and aCAPs before (blue) and after (red) AAV-ChR2(H134R) injeciton. B, the ABR threshold of AAV-injeciton(n=20) or saline-injeciton (n=20) after 4-6 weeks.C, the latent period of ABR before and after virus injection.D, oCAPs of a ChR2(H134R) guinea pig before and after laser stimulus:470nm blue light power laser;5.80mW/mm2 at the indicated duration at 5 Hz; 50 trials (oCAP-control:before stimulus;oCAP-10:after stimulus) .E, Representative aCAP in response to stimulation with 20 Hz average of 1024 trials).Representative oCAP in response to 470nm bule laser stimulation at 3.70mW and 11.1 Hz (average of 100 trials).F,the average amplitudes of oCAPs from a representative AAV-ChR2 (H134R)–injected adult guinea pig with the increasing of the laser intensities (from 1 to 10 level).

## Discussion

Here, guinea pig was established for developing optogenetic stimulation of the cochlea expressing ChR2 as an adult rodent model. The AAVs as the vectors needed for neural transduction should be optimized in order to safely and efficiently deliver the ChR2 (H134R), since they enable the foreign protein to be expressed at high rates and over long periods of time^[17].^Using AAV-ChR2 (H134R) vectors bearing the relatively short hSyn I promoter, we achieved neuron-specific transgene expression on the membrane of SGNs of guinea pigs. We found a more efficient and safer methods for improved transduction of virus-injection to express ChR2 in adult guinea pig. The injection of AAV-ChR2 (H134R) though round window achieved ChR2 (H134R) expression in about 90% of the injected cochlea, which contrasts the about 50% obtained with intra-modiolar injection^[16].^.Current studies not only improved transduction rates across all through the cochlea in the injected ear but also did not cause obvious SGN-loss for the vast majority of injected mice^[18, 19]^. A critical question about the process of virus injection through round window and expression of ChR2 (H134R) in SGNs is whether this caused any detrimental effect on auditory function in adult guinea pigs. We found the hearing results of average aABRs and aCAPs before and after the AAV-ChR2(H134R) injection. Both suggests that there is no significant detriment to the auditory pathway caused by virus-injection and ChR2 (H134R) expression. Several ways of AAV vectors injection have been proved that any obvious signs of neuronal loss or acoustic ABR decrease are caused in the cochlea^[9, 20]^. A significant advantage of optogenetic technology is the ability to control SGNs expressing ChR2 (H134R). To confirm expression of ChR2 (H134R) in adult guinea pig, we test the expression at both molecular and cellular level. Immunohistochemical and RT-PCR analysis revealed ChR2 (H134R) expression predominantly in the membrane of neurites and somata of SGNs.

Immunohistochemistry and colocalization suggested broad membrane expression of ChR2, which supported optogenetic high transduction rates in auditory SGNs. And there is no enough evidence for the ChR2 (H134R) expression beyond the injected ear. The evidences of establishing optogenetic models are highly relevant for experimental and clinical hearing research. The method above is a successful demonstration of adult cochlear transfection. Here, oCAPs and aCAPs were more similar in amplitude and showed the expected semblable latency for direct optogenetic SGN stimulation. Any considerable differences were found in the amplitudes and latent periods of oCAP among different experimental guinea pigs. The mild variability might be due to the efficient transduction rate, enough amount of ChR2 (H134R) expression among the transduced SGNs, and stable positioning of the optical fiber. Besides, we also explored various optical stimulation intensity. In 90% (18 of 20) of the tested animals, oCAP amplitude increases with light intensity. Considering the maximum light intensity safely used in our experiments(5.8 mW), which is within the safe range for optogenetic in vivo applications (up to~300 mW/mm2)^[21]^. Finally, the results of oCAPs were characterized by TDT recordings. Optical stimulation offers unprecedented opportunities to improve auditory prosthetics, because the adequate independent stimulation channels could enhance the coding frequency. Our study lays the groundwork for advancing optogenetics adult models for auditory research and future clinical applications. Such efforts include increasing transduction and expression rates by optimizing virus vectors and virus gene sequence, and better positioning of emitter in cochlea. The results of our present study demonstrate the feasibility of optogenetic activation of the auditory system after AAV-ChR2 (H134R) transduction in SGNs in adult guinea pigs. Those data paved cochlear optogenetics as a potential hearing restoration technology and laid the foundation for in-depth clinical research.

## Acknowledgement

This work was supported by the active health project of the Ministry of Science and Technology (2020YFC2004001), NSFC grant (81470700) and the Municipal-School Cooperation Project of Nanchong Science and Technology Bureau (19SXHZ0417)

## Author contributions

NY designed experiments. CL, FS, and NY performed experiments. CL, FS, and NY analyzed data. CL and NY wrote paper.

